# Design-Build-Test-Learn guided engineering of a whole-cell pyruvate biosensor based on transcription factor

**DOI:** 10.1101/2025.07.14.664704

**Authors:** Zihan Gao, Maria Suarez-Diez, Pieter Candry

## Abstract

Whole-cell biosensors are powerful tools for metabolite monitoring, yet challenges such as narrow dynamic range and high leaky expression limit their broader applications. Here, we present a systematic workflow based on two Design-Build-Test-Learn (DBTL) cycles to develop and optimize a transcription factor-based pyruvate biosensor in *Escherichia coli*. In the first iteration of the cycle, we constructed a biosensor that responded to intracellular pyruvate levels within 0.05–10mM range. In the second cycle, we implemented design of experiment (DoE) to systematically explore combinatorial effects of promoters and ribosome binding sites (RBSs). A first set of experiments were designed to identify factors with a significant effect on biosensor performance. The results showed RBS of report gene significantly influenced dynamic range by modulating basal and maximum expression, while RBS of transcription factor affected signal span. The Akaike Information Criterion was used to select a model incorporating two main effects and one interaction effects. The best-performing strain exhibited an 18.5-fold increase in dynamic range and a 37.2-fold reduction in leaky expression. Quantification of intracellular pyruvate confirmed an operational range of 1.23–6.81 μmol/g DCW. Our work demonstrates the power of DBTL cycles with statistical modelling for biosensor engineering, enabling more precise metabolic regulation and screening applications.

## 1. Introduction

Whole-cell biosensors have been drawing increasing attention over the last few decades. They couple a biological recognition element with a transducer that translates the biorecognition event into measurable signals^1^. In contrast to traditional physical and chemical sensors, whole-cell biosensors are highly sensitivity, easy-to-manufacture and cost-effective^2, 3^, showing widespread applications in metabolic engineering^4–7^, disease diagnosis^8^, food contaminant detection^9^, and environmental monitoring^10^. Despite these advantages, their sensing performance often falls short of real-world detection requirements, particularly regarding restricted dynamic ranges, high leaky expression and limit of detection^11, 12^. Figure 1A conceptually illustrates these key characteristics that often require optimization.

Recent advances in synthetic biology have provided tools to improve biosensor performance, such as promoter and ribosome-binding site (RBS) engineering for precise control of gene expression^13, 14^. The performance can be optimized sequentially by tuning individual genetic factors in isolation^4, 15, 16^. However, the cost of constructing and testing large numbers of genetic variants limits our ability to identify high-performance designs^17^. Combinatorial optimization of circuits, based on the simultaneous optimization of multiple genetic factors facilitates biosensor development ^18^.

Design of Experiments (DoE) procedures are essential to probe multidimensional experimental space with minimum number of experimental runs (Figure 1B). Statistical model-based DoE procedures have been widely used to simultaneously optimize multiple factors in biological processes, including optimization of conditions^19^, design of the bioprocess^20^ or metabolic pathway regulation^21^. The process often starts by considering two levels (low and high) per factor. The experimental results are summarized in a polynomial model in which main effects (MEs) describe how the response changes when each factor is varied individually, while two-factor interactions (2FI) and multiple factor interactions (xFI) describe the effect of the simultaneous change of two or more factors. The significance of each model parameter can then be evaluated using analysis of variance (ANOVA). Exploring all possible combinations of factor levels can result in a large experimental burden as 2*^n^* experiments are needed for *n* factors and 2 levels. Instead, fractional factorial design is an efficient screening method to identify MEs with less experiments^22^(Figure 1C). Different factorial designs exist depending on the number of experiments, but may result in confounded 2FI and xFI^18^. For instance, resolution IV designs can identify MEs but confound 2FI among each other. Consequently, this approach can indicate if 2FI are important but cannot distinguish which 2FI is most important. Although 2FI are confounded, these designs allow weighing their importance compared to the individual effects^23^. Additionally, model selection criteria such as the Akaike Information Criterion (AIC) can further support model evaluation and refinement^24^. In this study, we set out to test whether integrating DoE strategies with DBTL cycles could enable efficient and systematic optimization of biosensor performance.

Here, we applied a Design-Build-Test-Learn (DBTL) cycle to guide the engineering of whole-cell biosensors, which offers a systematic framework to iterate the steps of genetic circuits design, circuit construction, high throughput screening, and statistical modeling^25–27^. This approach was applied to pyruvate biosensors as a case study, as pyruvate is an important node linking glycolysis to the tricarboxylic acid (TCA) cycle^28^ as well as towards the synthesis of amino acids such as threonine, arginine, and succinate^29, 30^. Traditional analytical methods for pyruvate (e.g., HPLC, fluorimetric assays) are cumbersome and incapable of quantitative monitoring in living cells^25, 31^. Pyruvate biosensors would allow measuring intracellular pyruvate levels and dynamically control bioprocesses^7, 32, 33^. Building on these advancements, we implemented a two-cycle DBTL workflow to engineer a pyruvate-responsive biosensor in *Escherichia coli*. In the first cycle, we constructed a PdhR-based pyruvate biosensor in *E.coli ΔpdhR* strain. Then, in the second cycle, a Resolution IV fractional factorial design was used to explore the effect of two promoters and two RBSs on biosensor performance. A statistical model then supported the selection of the optimized biosensor design, which was used to validate the relationship between intracellular pyruvate concentration and biosensor output. Overall, the statistical DoE methodology proved to be a powerful and efficient strategy for enhancing biosensor performance and establishes a platform for intracellular pyruvate monitoring.

**Figure 1.**
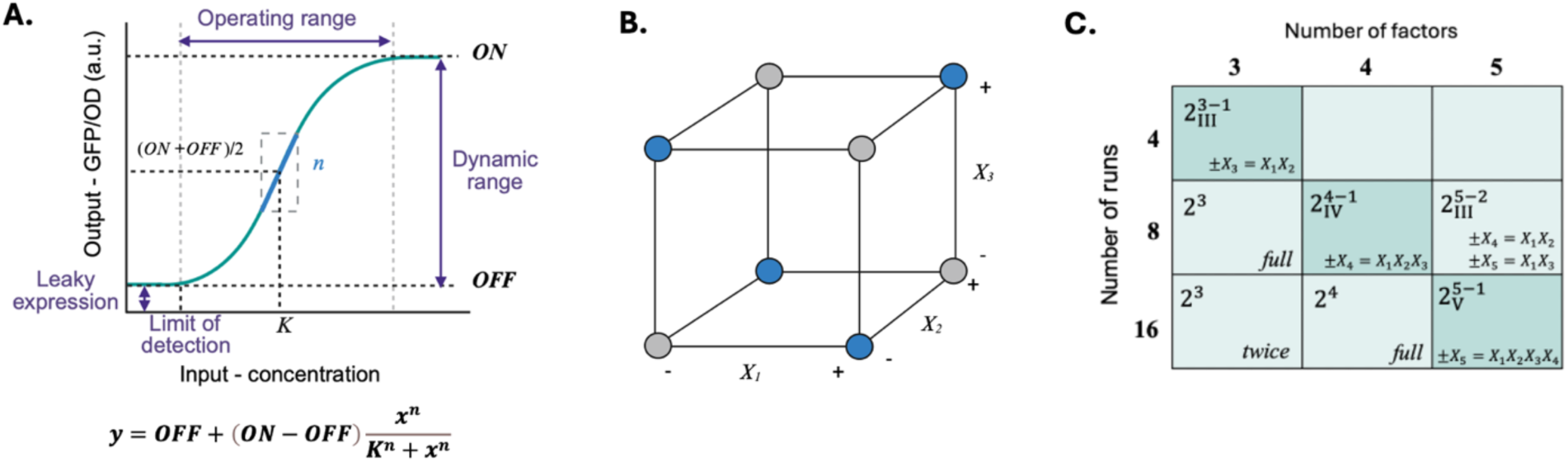
Overview of biosensor response modeling and design of experiments (DoE) strategy. (A) The dose-response model is fitted to a Hill’s equation. *Dynamic range* is defined as the ratio of maximum output signal (ON state) to the minimum output signal (OFF state). *Leaky expression* refers to the output level in the absence of input signal (OFF state). The *limit of detection* (LOD) is the lowest concentration of input which can be detected from background response. *Operating range* is the detectable concentration range of target concentration. These characteristics are commonly used to define the response of biosensor. (B) Schematic representation of the design space in a factorial DoE framework. (C) Fractional factorial design matrix illustrating the trade-off between the number of factors, resolution, and experimental runs. Higher resolution designs ensure the identification of MEs and interactions at the cost of increased experimental workload. Resolution III designs require fewer experiments but cannot reliably distinguish MEs from 2FI. Resolution IV designs allow the identification of MEs while confounding 2FI with one another. Resolution V designs allow identification of MEs and 2FI while confounding 3FI among each other^34^.

## 2. Results

To systematically to develop and optimize the performance of a whole-cell pyruvate biosensor, we implemented two rounds of DBTL cycle in *E. coli* ΔpdhR. The following sections describe the construction, testing, and optimization steps. Figure 2 illustrates the workflow of both DBTL cycles. In the first cycle, a PdhR-based genetic circuit was constructed to detect intracellular pyruvate within 0.05 – 10 mM. In the second cycle, a fractional factorial design guided optimization, leading to an improved biosensor with enhanced 18.54-fold dynamic range.

**Figure 2.**
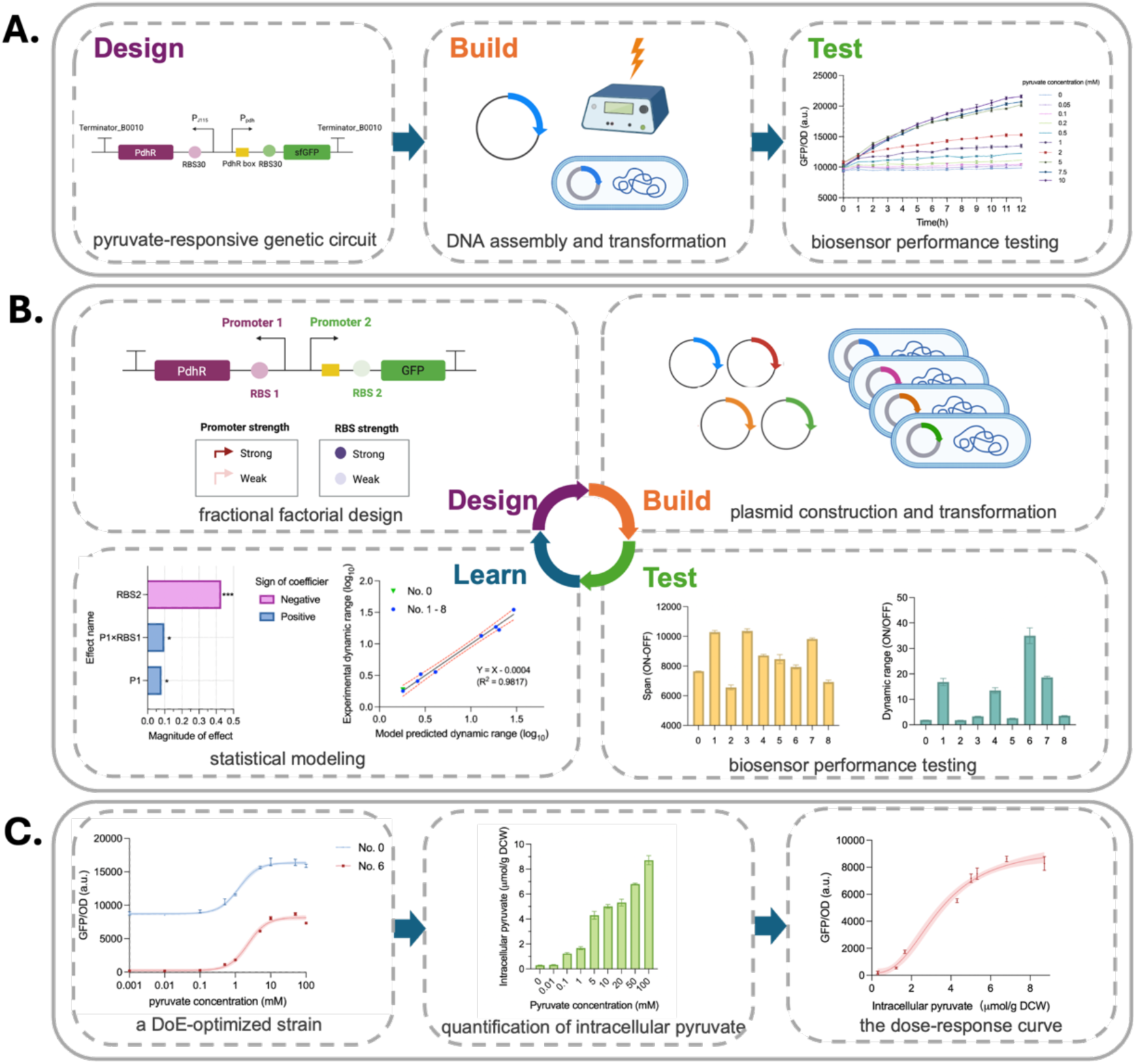
Overview of two rounds of DBTL cycle for the development and optimization of the pyruvate biosensor. (A) In the first DBT cycle, we designed a PdhR-based genetic circuit to detect intracellular pyruvate concentration. The genetic circuit was assembled into plasmid and transformed into *E.coli* Δ*pdhR*, and biosensor performance was evaluated. (B) In the second DBTL cycle, a fractional factorial Design was used to combine two promoters and two RBSs. These designs were then Built by transforming assembled plasmids into *E.coli* Δ*pdhR*. The performance of the transformed strains was Tested by fitting biosensor outputs to a Hill curve to evaluate biosensors’ span (ON-OFF) and dynamic range (ON/OFF). In the Learn stage, statistical modeling revealed main effects influencing dynamic range, guiding the selection of an optimized strain. (C) Finally, the optimized strain was validated to show improved performance, and the response was linked to intracellular pyruvate concentrations.

### 2.1 DBT Cycle 1: Construction of a PdhR-based pyruvate biosensor

In the first round of DBT, we constructed a PdhR-based biosensor for detecting intracellular pyruvate in *E. coli* Δ*pdhR*.

#### 2.1.1 Design: pyruvate-responsive genetic circuit

As a first step, the biosensor design leveraged PdhR as a pyruvate-sensing repressor to control *sfGFP* expression as a function of intracellular pyruvate concentration (Figure 3A). The gene circuit consisted of PdhR from *E.coli* MG1655 under the control of the constitutive promoter P_J115_. Then, the reporter gene *sfGFP* was selected for its enhanced folding efficiency and brightness, enabling accurate and robust quantification of gene expression in aerobic environment. *sfGFP* expression was driven by the Ppdh promoter, which contains the PdhR binding site. Both genes were equipped with ribosome binding sites (RBS30) to ensure proper translation efficiency (Figure 3B).

#### 2.1.2 Build: DNA assembly and transformation

In the build stage, the genetic circuit was assembled by HiFi DNA assembly into the pUC19 plasmid. Since the *E. coli* genome contains the *pdhR* gene, we chose to use a *pdhR* knockout strain to eliminate the influence of genomic *pdhR* on the experiment. Transformed strains were verified by colony PCR and Sanger sequencing.

#### 2.1.3 Test: biosensor performance testing

In the presence of 0.05–10 mM pyruvate, the relative fluorescence intensity of the biosensor increased with rising pyruvate concentrations (Figure 3C). This indicated that pyruvate effectively activated the biosensor. The 6-hour time point was selected as it provided a distinguishable GFP/OD signal across pyruvate concentrations, while also minimizing the need for extended incubation. At this time, the biosensor exhibited a linear response within the 0.1–5 mM range (R² = 0.97), which supported its application for quantitative pyruvate detection within this range(Figure 3D). Last, based on the significant difference in GFP/OD signal between 0.05 mM and 0 mM pyruvate (*P* < 0.001), the limit of detection was placed at 0.05 mM (Figure 3E).

**Figure 3.**
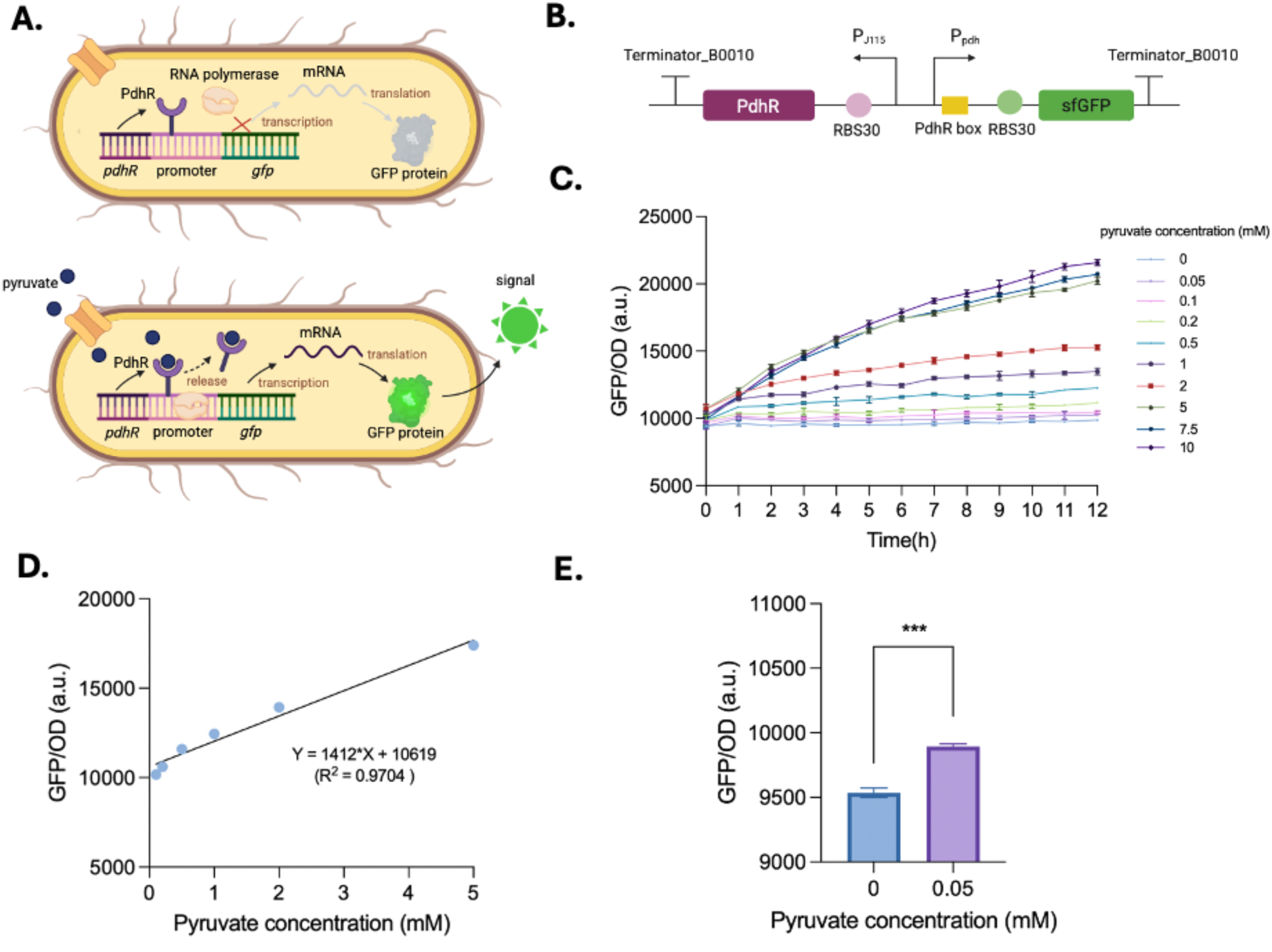
The construction and characterization of PdhR-based pyruvate biosensor. (A) Schematic representation of PdhR-based biosensor. In the absence of pyruvate, PdhR can recognize and bind to the PdhR box on the promoter of *sfGFP*, repressing its transcription and preventing *sfGFP* expression^35^. When pyruvate is present, it binds to PdhR, inducing a conformational change, which releases the promoter that can be bound to the RNA polymerase to carry out *sfGFP* transcription and translation. Within a certain range, the fluorescence intensity of sfGFP reflects the intracellular concentration of pyruvate. (B) Genetic circuit of PdhR-based biosensor. A constitutive promoter P_J105_ and RBS30 controls the expression of PdhR, which binds to PdhR box in P_pdh_. (C) The dose dependence of biosensor response to pyruvate. Measurements indicate mean and standard deviation of triplicate experiments. (D) The linear relationship between pyruvate concentration and GFP/OD at 6 h. (E) Statistically significant response to 0.05 mM pyruvate at 6 h, *** indicated *P* < 0.001.

### 2.2 Design and validation of a PdhR-regulated hybrid promoter

To assess the modularity and functionality of regulatory factors within the biosensor framework, we designed a hybrid promoter, P_J106_PdhR box_ and used this to control *sfGFP* expression (Figure 4A). To further investigate the regulatory role of PdhR, we compared GFP in the PdhR-knockout strain to a strain containing PdhR (Figure 4B). If the hybrid promotor functions as intended, GFP transcription would be repressed by PdhR, leading to a higher GFP signal in a non-PdhR-strain compared to a PdhR-containing strain. At 0 h, the strain without PdhR showed significantly higher GFP expression than the strain with PdhR (Figure 4C; *P* < 0.001). This indicated the inhibitory effect of PdhR already existed during the LB growth stage. When the strains were then cultivated in M9 medium containing 10 mM pyruvate, both groups showed increased GFP/OD values compared to 0h, but the strain without PdhR remained significantly higher than the strain with PdhR (*P* < 0.001). Additionally, the strain with PdhR demonstrated a significant increase in GFP/OD following pyruvate treatment, confirming that pyruvate can activate the biosensor by relieving PdhR-regulated repression. These results illustrate PdhR can inhibit the transcription of the hybrid promoter P_J106_PdhR box_. This validation step ensured that the engineered promoter retained functionality for the following combinatorial design experiments.

**Figure 4.**
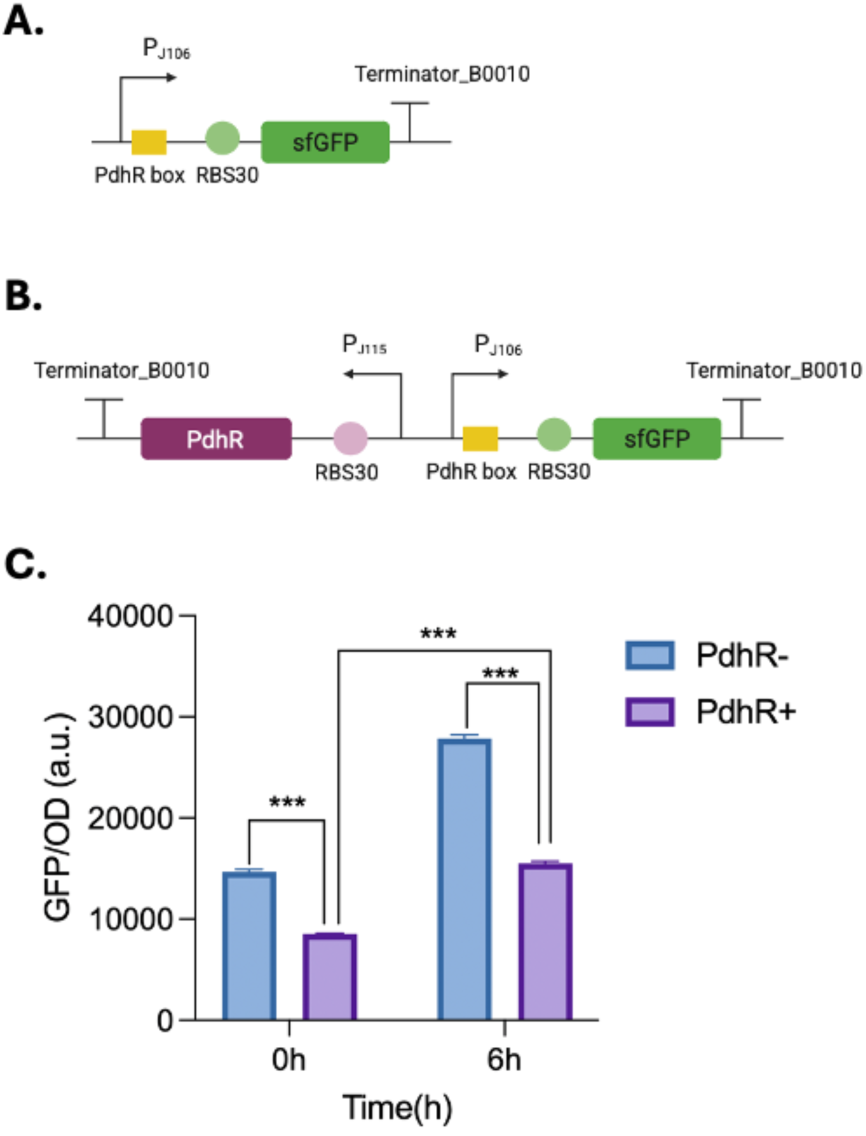
The construction and functional validation of hybrid promoter. (A) Schematic diagram of integration of PdhR box into constitutive promoter P_J106_. (B) Schematic diagram of hybrid promoter in strain with PdhR. (C) The strength of hybrid promoter in strains with (purple column) or without PdhR (blue column) at 0 h and 6 h, *** indicated *P* < 0.001.

### 2.3 DBTL Cycle2: DoE to optimize performance

To optimize the performance of the biosensor, we aimed to simultaneously maximize its dynamic range (ON/OFF), minimize background expression (OFF) and increasing span (ON-OFF). This could be achieved through systematic combinations of promoters and RBSs at different strength guided by fractional factorial design. Our goal was to develop a statistical model that can screen the main factors and interactions, providing valuable insights for future efforts in biosensor construction, optimization, and fine-tuning.

#### 2.3.1 Design: fractional factorial design

In our study, we conducted a round of DoE to optimize the performance of the PdhR-based biosensor. We focused on four key factors: (i) P1: the promoter of PdhR, which regulates the transcription of PdhR, (ii) RBS1: the ribosome binding site of PdhR, which modulates the translation of PdhR, (iii) P2: the promoter of sfGFP, which controls the transcription of report gene, and (iv) RBS2: the ribosome binding site of sfGFP, which determines the translation level of sfGFP). Each factor was assigned discrete levels: the low level coded as −1, and the high level as 1. Given the size of the screened library, a total of 16 combinations would be needed to fully explore the combinatorial space. We selected a fractional factorial design to reduce the experimental effort. Here a resolution IV design was implemented as it can identify the main effects and whether two-factor interactions affect biosensor performance. In our case, a resolution IV design with 4 factors resulted in 2^4–1^ = 8 experiments (strains), as illustrated in Figure 1C.

#### 2.3.2 Build: plasmid construction and transformation

We assembled eight genetic circuits with different promoters and RBSs, driven by strong (1) or weak (-1). The No. 0 was regarded as control, with P2 in this strain being P_pdh_ from *E.coli* genome (Table 1, S5).

**Table 1.** Fractional factorial design to screen factors that affect biosensor performance.

| No. | P1 | RBS1 | P2 | RBS2 | OFF (a.u.) | ON (a.u.) | Span (ON-OFF) | Dynamic range (ON/OFF) | OFF ($\log_{10}$) | ON ($\log_{10}$) | Dynamic range ($\log_{10}$) |
| --- | --- | --- | --- | --- | --- | --- | --- | --- | --- | --- | --- |
| 0 | -1 | 1 | n.a | 1 | $8720.7 \pm 24.2$ | $16363.0 \pm 25.4$ | $7642.7 \pm 26.1$ | $1.88 \pm 0.01$ | $3.9405 \pm 0.0012$ | $4.2139 \pm 0.0007$ | $0.2733 \pm 0.0011$ |
| 1 | -1 | -1 | -1 | -1 | $654.0 \pm 54.5$ | $10937.0 \pm 59.6$ | $10283.0 \pm 113.8$ | $16.72 \pm 1.42$ | $2.8156 \pm 0.0354$ | $4.0389 \pm 0.0024$ | $1.2233 \pm 0.0378$ |
| 2 | -1 | 1 | -1 | 1 | $8286.7 \pm 107.6$ | $14841.0 \pm 144.4$ | $6554.3 \pm 167.0$ | $1.79 \pm 0.03$ | $3.9184 \pm 0.0056$ | $4.1715 \pm 0.0042$ | $0.2531 \pm 0.0066$ |
| 3 | -1 | -1 | 1 | 1 | $4495.0 \pm 95.6$ | $14857.7 \pm 59.5$ | $10362.3 \pm 145.6$ | $3.31 \pm 0.08$ | $3.6527 \pm 0.0092$ | $4.1720 \pm 0.0017$ | $0.5192 \pm 0.0106$ |
| 4 | -1 | 1 | 1 | -1 | $700.8 \pm 58.4$ | $9408.7 \pm 33.2$ | $8708.0 \pm 82.3$ | $13.43 \pm 1.10$ | $2.8456 \pm 0.0354$ | $3.9735 \pm 0.0015$ | $1.1279 \pm 0.0363$ |
| 5 | 1 | -1 | -1 | 1 | $5415.0 \pm 105.3$ | $13882.0 \pm 199.9$ | $8466.7 \pm 301.0$ | $2.56 \pm 0.09$ | $3.7336 \pm 0.0085$ | $4.1425 \pm 0.0062$ | $0.4089 \pm 0.0145$ |
| 6 | 1 | 1 | -1 | -1 | $234.3 \pm 16.6$ | $8168.7 \pm 124.9$ | $7934.3 \pm 140.6$ | $34.86 \pm 3.10$ | $2.3698 \pm 0.0314$ | $3.9122 \pm 0.0066$ | $1.5424 \pm 0.0375$ |
| 7 | 1 | -1 | 1 | -1 | $558.3 \pm 12.5$ | $10374.0 \pm 65.5$ | $9816.0 \pm 76.7$ | $18.58 \pm 0.52$ | $2.7469 \pm 0.0097$ | $4.0159 \pm 0.0027$ | $1.2691 \pm 0.0122$ |
| 8 | 1 | 1 | 1 | 1 | $2687.0 \pm 33.5$ | $9612.7 \pm 129.8$ | $6926.0 \pm 124.7$ | $3.58 \pm 0.06$ | $3.4293 \pm 0.0054$ | $3.9828 \pm 0.0059$ | $0.5536 \pm 0.0068$ |

<sup>a</sup> OFF and ON are the result of the fit to the Hill equation (see Materials & methods). The corresponding Hill coefficients are listed in Table S5. All the reported values indicate the mean
of three biological replicates with ± denoting the standard deviation (SD) of three replicates. n.a represented not applicable.

#### 2.3.3 Test: biosensor performance testing

We assessed the performance of the pyruvate biosensors by measuring GFP/OD at various pyruvate concentrations and fitting the data to a Hill curve. The OFF state corresponds to the minimum expression of the biosensor, while the ON state represents the maximum expression. The biosensors’ span and dynamic range were calculated (Table 1).

#### 2.3.4 Learn: statistical modeling

The tested strains varied in performance of both span and dynamic range. The span varied up to 1.36-fold (Figure 5A). ANOVA analysis indicated that effects of P1, P2 and RBS2 were insignificant (*P* > 0.05). RBS1, on the other hand, had a significant negative effect on the span (*P* = 0.011, Figure 5B, 5C, Table S6). Dynamic ranges varied from 1.88 ± 0.01 to 34.86 ± 3.10 (a.u.) (Figure 5D). ANOVA analysis showed that RBS2 had a significant negative effect on the dynamic range (*P* = 0.0066), whereas the effects of other factors were not significant (*P* > 0.05) (Figure 5E, 5F, Table S6). The regression models for span and dynamic range exhibited good fit to the experimental data, with adjusted R² values of 0.85 and 0.86, and *P* values of 0.038 and 0.035, respectively, indicating statistical significance and joint contributions of the selected factors. These coefficients in the model indicated that RBS1 played a crucial role in modulating the span, while RBS2 strongly influenced the dynamic range. Both showed a negative effect.

**Figure 5.**
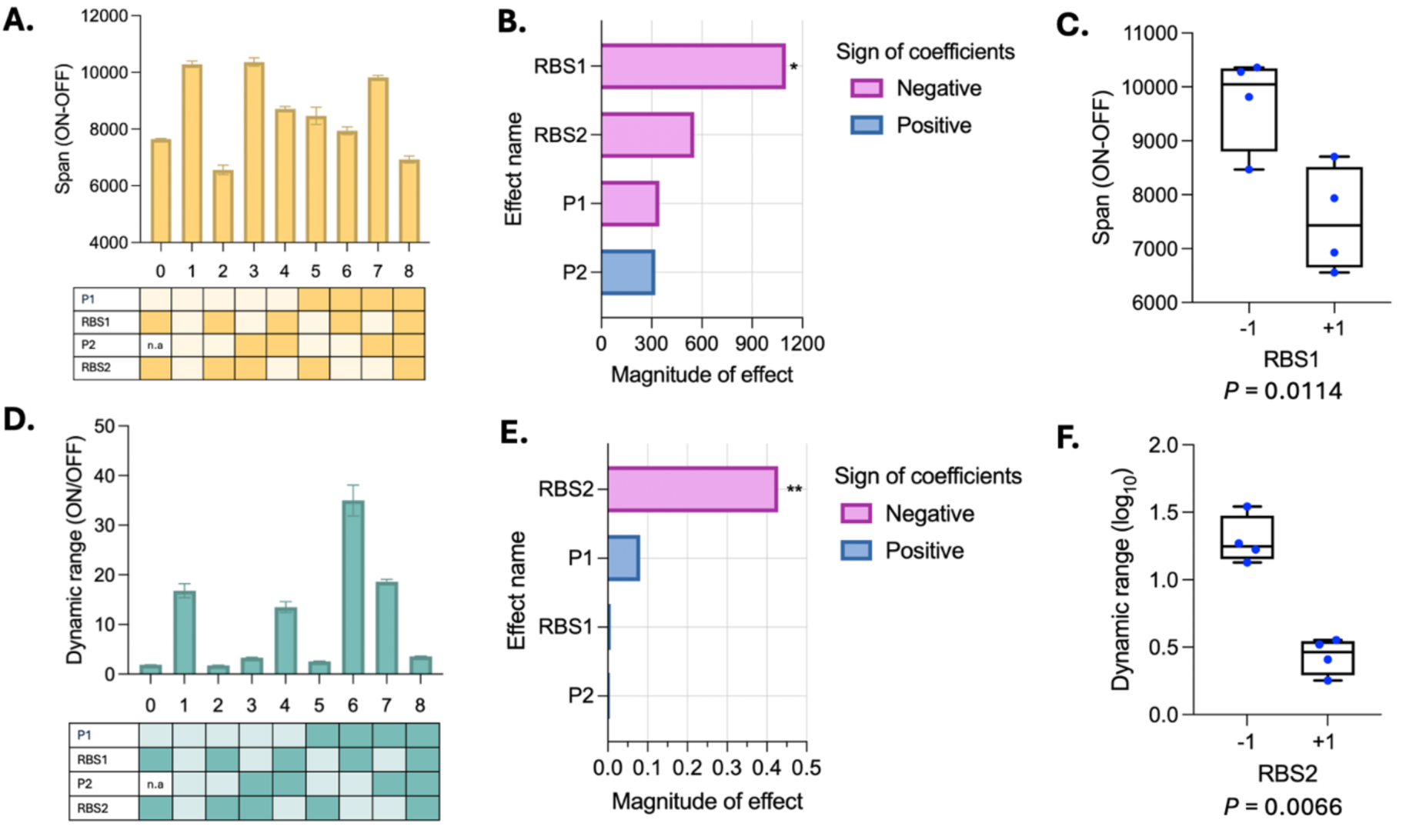
Screening and statistical analysis of main factors effecting biosensor performance. (A) The span of multiple tested biosensor variants . Light color indicated low level (-1) and dark color indicated high level (1). (B) Coefficients of model including main effects on span. The magnitude of effect refers to the value of the estimated coefficient from the linear mode * indicated *P* < 0.05, ** *P* <0.01. (C) ANOVA analysis of the effect of RBS1 on span. ANOVA analysis data were obtained from the fractional factorial design. The box displays the interquartile range (IQR), with the line indicating the median, and whiskers extending to the minimum and maximum values. (D) The dynamic range of multiple tested strains. (E) Coefficients of model including main effects on log_10_-transformed dynamic range. (F) ANOVA analysis of the effect of RBS2 on log_10_-transformed dynamic range.

Given that RBS2 negatively affects the dynamic range, we further investigated its impact on both OFF and ON states to understand the underlying mechanism. When a strong RBS2 was selected, it significantly impacted both the OFF and ON states, leading to increased leaky expression (Figure 6A, 6B, Table S6) and a higher maximum expression level (Figure 6C, 6D, Table S6). The effect of RBS2 on both the OFF and ON states was significant (*P* < 0.05), particularly in the OFF state, where its influence was more pronounced. The increased translation efficiency driven by RBS2 likely elevated basal expression levels, leading to leaky expression in the OFF state.

This leaky expression in the OFF state is particularly critical, as it directly limits the biosensor’s dynamic range by raising the signal baseline. Therefore, minimizing RBS2-driven translation in the OFF state is essential for maximizing dynamic range.

**Figure 6.**
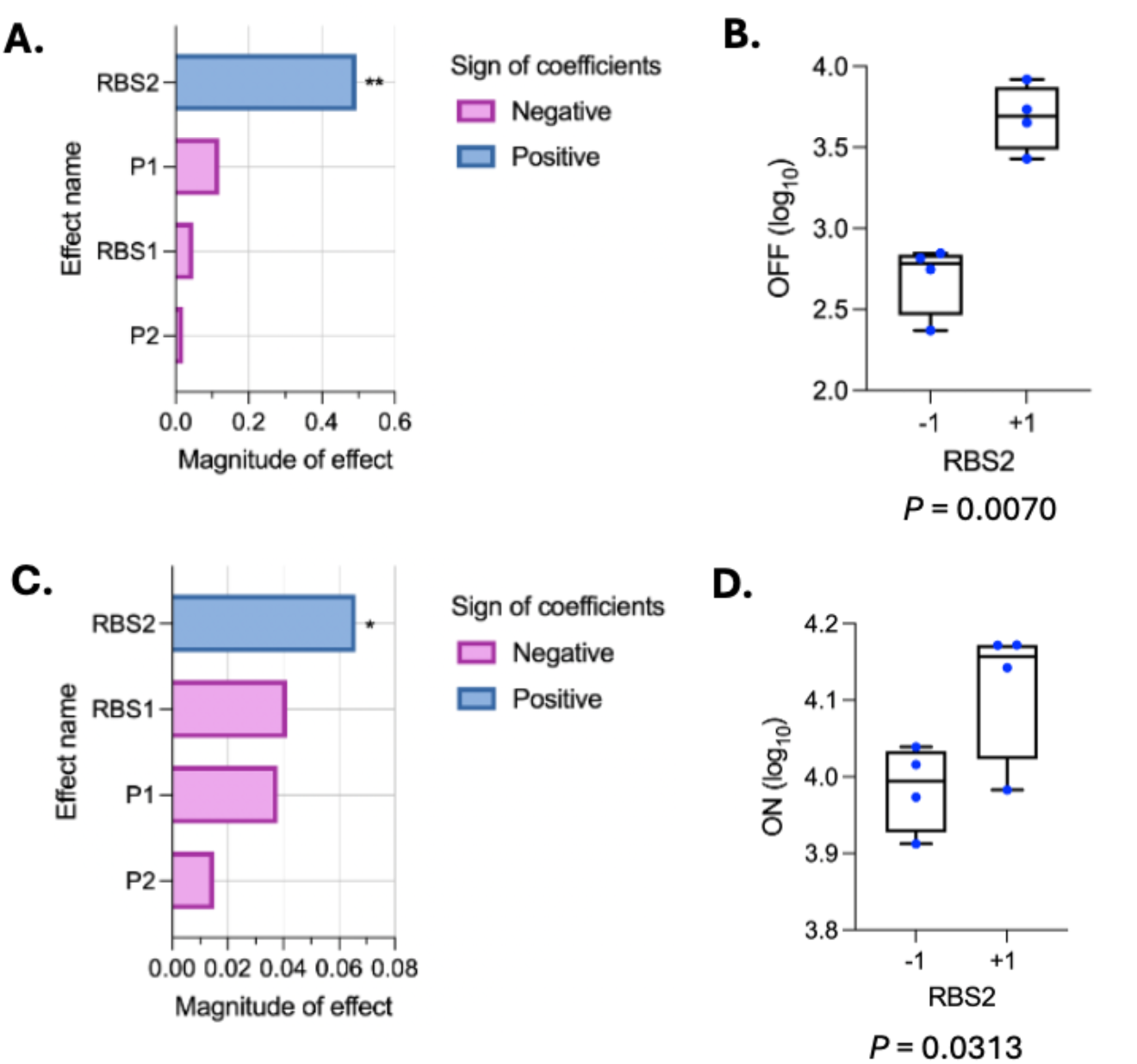
Statistical analysis of RBS2 effecting dynamic range. (A) Coefficients of model including main effects on log_10_-transformed OFF state, * indicated *P* < 0.05, ** *P* <0.01. (B) ANOVA analysis of the effect of RBS2 on log_10_-transformed OFF state. (C) Coefficients of model including main effects on log_10_-transformed ON state. (D) ANOVA analysis of the effect of RBS2 on log_10_-transformed ON state.

To further understand these effects, we analyzed all interactions between factors (Figure S1). The result showed the P1×RBS1 interaction had a greater effect magnitude compared to other interactions. In resolution IV design, the interaction terms P1×RBS1 and P2×RBS2 are aliased, meaning their effects cannot be distinguished statistically (Table S6). Then, we fitted the model using the four main effects and the P1×RBS1 interaction, which showed that RBS1 and P2 had no significant effect (*P* > 0.05) (Figure S2). We used the AIC for model comparison and Model 4 (*Y* = β_0_ + β_1_⋅*P1*+ β_2_⋅*RBS2* +β_3_⋅ *P1*⋅ *RBS1*) was selected as the best performing model (Table S7). In the model, we showed P1×RBS1 to represent this confounded 2FI. *Y* represent the predicted log_10_-transformed dynamic range. According to the final regression coefficients in the linear model, RBS2 negatively impacted the system (β_2_ = -0.43, *P* = 0.000075), while P1 and the P1×RBS1 interaction contributed positively to the dynamic range (β_1_ = 0.080, *P* = 0.034; β_3_ = 0.097, *P* = 0.019) (Table S6, Figure 7A). The effect of P1 on dynamic range was dependent on the level of RBS1 (Figure 7B). When RBS1 was at high level (1), the dynamic range increased with P1, indicating a positive correlation. At the low level of RBS1 (-1), the dynamic range slightly decreased as P1 increased, showing a weak negative correlation. This suggested that P1 had a notable effect on the dynamic range, and its impact was modulated by RBS1. The interaction of P2×RBS2 showed the effect of the promoter strength on the dynamic range depends on the level of RBS2 (Figure S3). The proposed regression model demonstrated a good fit with the experimental data, having an adjusted R^2^ of 0.98 (Table S6). The relationship between the model-predicted and experimentally observed dynamic ranges for variants No. 1-8 showed a linear relationship (R^2^ = 0.98). To further validate the model, strain No. 0 data were projected in formula and the result fell within the 95% confidence interval (CI) of the curve (Figure 7C), confirming the accuracy of the model. The linear regression model enabled us to determine the optimal gene expression levels for the genetic circuits’ factors, leading to variant No. 6 having 18.54-fold increase in the dynamic range and a 37.22-fold reduction in leaky expression. The Hill’s slope of variant No. 0 was 1.57, while that of variant No. 6 was 1.64, indicating a slightly higher response to pyruvate (Figure 7D). These results demonstrate the effectiveness of linear regression model as a systematic tool for guiding the rational design of regulatory factors to enhance biosensor performance.

**Figure 7.**
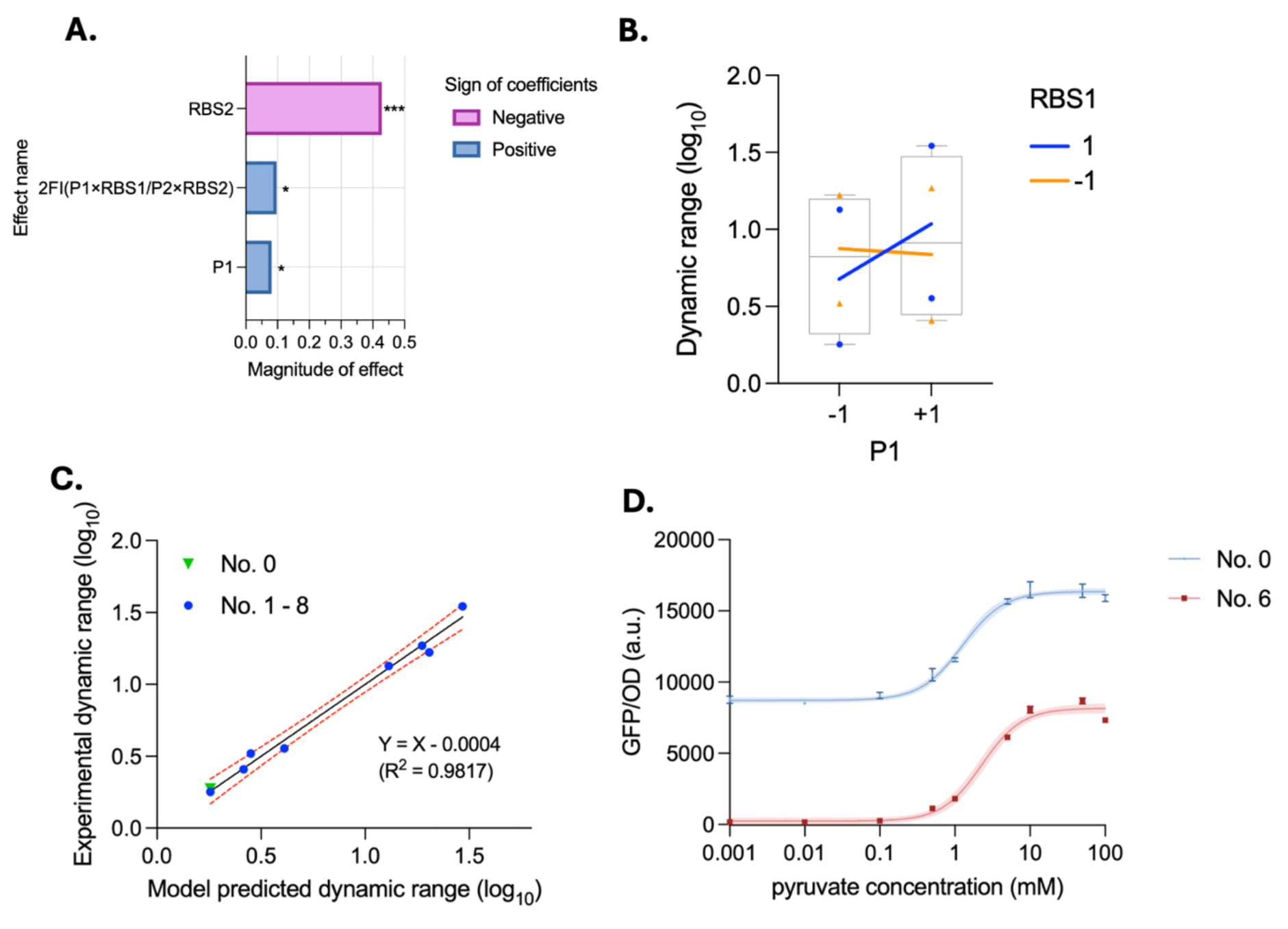
Summary of fractional factorial design results and validation of model prediction. (A) Linear regression analysis of Model 4 (*Y* = β_0_ + β_1_⋅*P1*+ β_2_⋅*RBS2* +β_3_⋅ *P1*⋅ *RBS1*), * indicated *P* < 0.05, \*\*\**P* < 0.001. (B) Interaction effects between P1 and RBS1. Blue circles and orange triangles indicate data points under high (1) and low (-1) levels of RBS1, respectively. The lines connect group means and illustrate the interaction effect: under high RBS1 (blue line), whereas under low RBS1 (orange line). (C) The variants No. 1-8 (blue dots) correlation between model predicted and experimentally measured dynamic ranges. The control No. 0 (green triangle) was in 95% CI area. (D) The dose–response curves of control No. 0 and optimized variant No. 6.

### 2.4 Quantification of pyruvate in a DoE-optimized variant

Based on our linear regression model, variant No. 6 exhibited the highest dynamic range and lowest leaky expression. To evaluate this variant’s practical applicability, we quantified extracellular and intracellular pyruvate concentration and linked this to biosensor output. Across a range of initial pyruvate concentrations (0-100 mM), we measured extracellular pyruvate concentrations at 0 h and 6 h (Figure 8A). At 0 h, the HPLC results confirmed that the measured extracellular concentrations matched the amounts added, down to the detection limit of 0.1 mM. When pyruvate was used as carbon source, the extracellular concentration decreased by approximately 20 mM after 6 hours, indicating pyruvate consumption. Intracellular pyruvate levels increased with rising extracellular concentrations, suggesting pyruvate transport into the cell (Figure 8B). Furthermore, by fitting GFP/OD-data as a function of intracellular pyruvate concentrations to the Hill equation, we determined the operational range of 1.23–6.81 μmol/g DCW. This indicated the biosensor had a high sensitivity to pyruvate (Figure 8C). Taken together, these results demonstrated that variant No. 6 is a promising candidate for pyruvate-responsive applications, combining tight gene regulation with high responsiveness.

**Figure 8.**
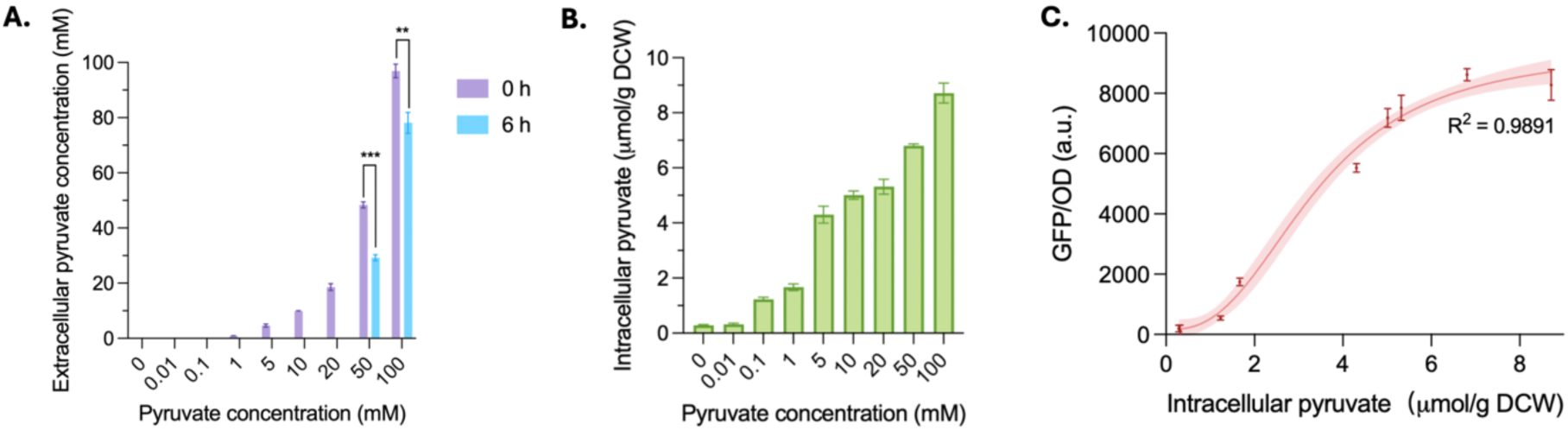
Quantification of extracellular and intracellular pyruvate concentration. (A) Extracellular pyruvate concentration at 0 h (purple column) and 6 h (blue column) after supplementation with 0, 0.01, 1, 5, 10, 20, 50, 100 mM pyruvate, ** indicated *P* < 0.01, \*\*\**P* < 0.001. (B) Intracellular pyruvate concentration measured at 6 h. (C) The dose–response curves of biosensor response to intracellular pyruvate. Data were fitted to a Hill equation (Table S8). Shaded area represents 95% CI.

## 3. Discussion

The Design-Build-Test-Learn (DBTL) cycle provides a systematic framework to develop and optimize whole-cell biosensors. In this study, we implemented two rounds of DBTL to optimize a PdhR-based pyruvate biosensor in *E. coli*. We systematically screened combinations of promoters and RBSs using a fractional factorial design and identified key factors affecting biosensor behavior. We developed a PdhR-based pyruvate biosensor which can respond to 0.05– 10 mM pyruvate and drastically increased the dynamic range by 18.54-fold and decreased leaky expression by 37.22-fold. The operational intracellular pyruvate range of the DoE-optimized variant was quantified as 1.23–6.81 μmol/g DCW. Previously developed PdhR-based biosensors developed by Wang et al. in *Candida glycerinogenes* responded to 18.2–272.5 mM pyruvate, with dynamic range at approximately 3.5, corresponding to intracellular concentration range of 0.9–4.2 μmol/g DCW^36^. Xu et al. also reported a pyruvate-responsive genetic circuit in *Bacillus subtilis* that responded to 45.5–181.8 mM pyruvate, with a dynamic range of approximately was4^32^. In comparison, the pyruvate biosensor developed in this study exhibited a broader dynamic range and higher sensitivity at lower extracellular pyruvate concentrations. These differences in response range and intracellular concentrations reflect host-specific differences in pyruvate metabolism, emphasizing the significance of choosing the appropriate chassis in the biosensor design and application.

To achieve this improved biosensor performance and characterize the factors contributing to its performance, we proposed resolution IV design as a suitable balance between information gain and experimental effort, making it a good option to screen the effect of factors in the response^23^. Our approach employed a four-factor, two-level fractional factorial design that successfully identified key regulatory factors and potential interactions. Compared to the three-factor, three-level optimization employed by Berepiki et al.^37^, our method provided greater efficiency with fewer experimental runs and demonstrated the power of statistical modeling in guiding biosensor engineering. Although biosensor responses are inherently nonlinear and follow a sigmoidal Hill curve, our work shows the possibility of fitting linear models to this behaviour by focusing on the Hill curve parameters. Expressing the dynamic range as the ratio of ON to OFF levels quantifies the fold change between ON and OFF, rather than characterizing the entire response curve. As such, it is possible for the ON/OFF ratio to be used as linear performance indicator. This approach allows effective comparison and optimization of biosensor performance, regardless of the specific shape of the dose–response curve.

The strength of RBSs and promoters effectively modulated biosensor output by balancing repression strength and reporter expression levels. The observed effects of RBSs can be attributed to their role in controlling translational efficiency, which in turn influences protein abundance and folding^15^. High expression of PdhR due to a strong RBS1 resulted in enhanced repression, reducing signal span. Similarly, RBS2 was found to significantly influence the dynamic range by modulating both basal (OFF) and maximum (ON) expression. When RBS2 was at high level, it led to a higher leaky expression as well as increased maximum expression, whilst a low translation rate reduce both leaky expression and also maximum expression^38^. Since dynamic range is defined as the ratio between maximum and leaky expression of the reporter, a weaker RBS2 lowered translation rate, which resulted in improved biosensor performance. This indicated that adjusting RBS strength was a simple yet effective strategy to modulate translation rates and thereby optimize biosensor behavior^39^. In addition, P1 significantly affected dynamic range, highlighting that transcription level control of the PdhR modulates repression strength and output variability. Future optimization efforts can be guided by different objectives. When the goal is to further optimize dynamic range within the current design space, the most significant factor RBS2 can be fixed at its optimal level (e.g, RBS2 at -1 level), while another significant factor P2 can be further fine-tuned (e.g, intermediate values between –1 and +1) to explore nonlinear responses. Alternatively, factors with the most important main effects could be further optimized by moving outside the initial design space. For example, a weaker RBS2 and a stronger P2 than currently implemented could be considered to explore whether dynamic range can be improved further. Overall, these results demonstrated rational tuning of genetic circuits can fine-tune biosensor performance, and statistical modelling can capture meaningful mechanistic relationships within a transcription factor-based genetic circuit.

The accumulation of endogenous pyruvate in LB medium may further activate the biosensor, preventing complete repression and thereby reducing the contrast between ON and OFF states. This highlights the need to evaluate sensor performance under more realistic conditions. Unlike defined media with precisely pyruvate concentration, real-world environments often contain background metabolites that may induce unintended activation, particularly in the presence of carbohydrates. Bi et. developed a single-cell pyruvate fluorescence resonance energy transfer (FRET) biosensor to measure dynamic fluctuations in metabolic network of *E.coli*, they found rapid conversion of glucose to pyruvate occurs on timescale of second. Moreover, this FRET biosensor can also respond to fructose, acetate and glycerol^40^. Yang et. first verified PdhR specificity *in vitro* when developing a pyruvate biosensor in *Saccharomyces cerevisiae*. Their biosensor demonstrated that glucose and fructose activate cytoplasmic pyruvate levels more effectively than direct exposure to pyruvate, and that they do so in a dose-dependent manner^33^. This suggested that the metabolic environment may play a crucial role in shaping pyruvate biosensor performance, by affecting their basal expression, response dynamic range and response speed.

This optimized biosensor could also provide a versatile platform for metabolic engineering. First, it can act as a dynamic valve for controlling pyruvate flux in synthetic pathways, improving yields of pyruvate-derived compounds such as succinate or alanine^30, 41^. Second, it can serve as a high-throughput screening tool to identify pyruvate-accumulating strains or regulatory mutations^42^. Finally, the modular structure of the biosensor allows for further customization, including the integration of AI-designed synthetic TFs to expand responsiveness to pyruvate analogs^43–45^.

Overall, this work demonstrates how systematic implementation of DBTL cycles guided by statistical modelling such as DoE and AIC, which could enhance biosensor performance and provides a foundation for broader application in synthetic biology circuit design and precision metabolic engineering.

## 4. Methods

### 4.1 Strains and media

All strains used in this study are listed in Supplementary Table S1. *E. coli* DH5α was used for cloning, DNA assembly and plasmid construction. *E.coli* BW25113 *ΔpdhR* was used for biosensor performance assays. Strains were cultured in Lysogeny broth (LB) medium (10 g/L tryptone, 10 g/L NaCl, and 5 g/L yeast extract) with antibiotics. Pyruvate biosensor performance was tested in M9 minimal medium (3.88 g/L K_2_HPO_4_, 1.63 g/L NaH_2_PO_4_, 2 g/L (NH_4_)SO_4_, 0.01 g/L EDTA, 0.1 g/L MgCl_2_·6H_2_O, 0.002 g/L ZnSO_4_·7H_2_O, 0.001 g/L CaCl_2_·2H_2_O, 0.005 g/L FeSO_4_·7H_2_O, 0.0002 g/L Na_2_MoO_4_·2H_2_O, 0.0002 g/L CuSO_4_·5H_2_O, 0.0004 g/L CoCl_2_·6H_2_O, 0.001 g/L MnCl_2_·2H_2_O) with ampicillin (100 μg/mL) and kanamycin (50 μg/mL). Sodium pyruvate stock solutions were made in MilliQ H_2_O and filtered (0.20 μm) before use.

### 4.2 Construction of biosensor and transformation in *E.coli* Δ*pdhR* strain.

All plasmids used or constructed in this study were listed in Table S2, all genetic parts, sequences and primers were listed in Table S3 and S4. The *pdhR* gene and its promoter P_pdh_ used in the PdhR-based pyruvate biosensor were derived from *E. coli* MG1655. The *pdhR* gene was cloned from *E.coli* genome by Q5 polymerase (NEB, #M0491S). The pUC19 plasmid containing the ColE origin and ampicillin resistance marker was used as the vector. The constitutive promoter of *pdhR* gene and RBSs were obtained from the iGEM Parts Registry (https://parts.igem.org/Promoters). DNA oligonucleotides and fragments containing promoters and RBS were synthesized by IDT. Fragments were assembled into pUC19 via HiFi DNA Assembly (NEB, #E2621) following manufacturer’s protocols. PCR products were treated with DpnI (NEB, # R0176S) to remove methylated template DNA following manufacturer’s protocols. All constructs were verified by colony PCR with Phire II polymerase (Thermo Fisher, #F126S) following manufacturer’s protocols. PCR positive clones were subsequently validated by Sanger sequencing. Then, electroporation was used to introduce plasmids containing the designed genetic circuits into the KEIO-collection, kanamycin-resistant Δ*pdhR* strain^46^ (Table S1). Cells were electroporated using a Gene Pulser Xcell electroporator (Bio-Rad) following bacterial standard pre-set protocols (2.5kV, 0.1 cm cuvette). Following electroporation, cells were recovered in SOC medium (Super Optimal broth with Catabolite repression, 20 g/L tryptone, 5 g/L yeast extract, 0.584 g/L NaCl, 0.186 g/L KCl, 2.033 g/L MgCl_2_·6H_2_O, 2.465 g/L MgSO_4_·7H_2_O, and 3.603 g/L glucose 3.603 g/L) at 37°C for 1 hour to allow for plasmid expression. Transformants were then selected on LB agar plates containing kanamycin (50 μg/mL) and ampicillin (100 μg/mL) to ensure the presence of both the genomic kanamycin resistance gene and the ampicillin resistant plasmid. Transformants were screened by colony PCR using two sets of primers: one set of primers (verify_PG_fwd and verify_PG_rev) to verify correct construct integration of *pdhR* and *sfGFP* in plasmids and another set of primers (verify_PdhR_fwd and verify_PdhR_rev) to confirm *pdhR* gene knockout in *E.coli* Δ*pdhR* strain. Positive clones were further validated by Sanger sequencing and stored at -70°C in glycerol stocks.

### 4.3 Biosensor performance testing

Biosensor performance was tested by evaluating GFP production in response to varying pyruvate concentrations. Bacterial strains were streaked from glycerol stocks onto LB agar plates containing ampicillin (100 µg/mL) and kanamycin (50 µg/mL) and incubated overnight at 37°C. Colonies were inoculated into LB medium with antibiotics and cultured overnight at 37°C with shaking at 250 rpm. The following morning, 0.5 mL of the overnight culture was diluted into 4.5 mL of fresh LB medium with the antibiotics and grown for 6 hours to control background expression. Cells were washed twice with M9 minimal medium and cultured overnight in M9 with antibiotics. After washing once with M9, cultures were diluted to an initial OD₆₀₀ of 0.3.

Pyruvate was added at final concentrations of 0, 0.001, 0.01, 0.1, 0.5, 1,5,10,50,100 mM. A total of 150 µL of the culture was transferred to each well of a 96-well plate, covered with 50 µL of mineral oil to prevent evaporation, and incubated at 37°C with shaking at 731 cpm (2 mm orbit) for 6 hours. Fluorescence intensity (sfGFP, excitation at 485 nm; emission at 530 nm) and optical density (OD₆₀₀) were measured using a microplate reader (BioTek Synergy H1). Biomass growth was accounted for by normalizing GFP expression to OD₆₀₀. Background fluorescence of the strain was corrected by subtracting the GFP/OD values of the negative control (*E. coli ΔpdhR* strain).

### 4.4 Biosensor performance optimization

For biosensor performance optimization, we used the fractional factorial design of Resolution IV. Four factors were considered: (i) P1: a constitutive promoter of PdhR, (ii) RBS1: a ribosome binding site controlling the translation efficiency of PdhR, (iii) P2: a hybrid promoter controlling sfGFP transcription, and (iv) RBS2: a ribosome binding site regulating the translation efficiency of sfGFP. Each factor was tested at two levels: a high level (coded as +1) and a low level (coded as –1), resulting in a design with 2⁴⁻¹ = 8 experimental runs. The promoter of PdhR, a strong P_J108_ functioned as 1, while a weak P_J115_ acted as -1. For the hybrid promoter of GFP, the strong P_J106_PdhR box_ was 1, and the weak P_J114_PdhR box_ was -1. The hybrid promoters contained the PdhR box sequence ATTGGTAAGACCAAT in the core region of the constitutive promoter^47^. The distance between the PdhR box and the transcription start site “+1” was 11 base pairs^32^. Regarding RBS strength, the strong RBS30 was 1, whereas the weak RBS33 was -1. The constitutive promoters and RBSs were obtained from the iGEM Parts Registry. The fragments containing promoters and RBS were synthesized by IDT and assembled into pUC19 vector by using HiFi DNA Assembly following manufacturer’s protocols. The genetic configuration of constructed circuits and levels of four independent factors are listed in Figure 9.

**Figure 9.**
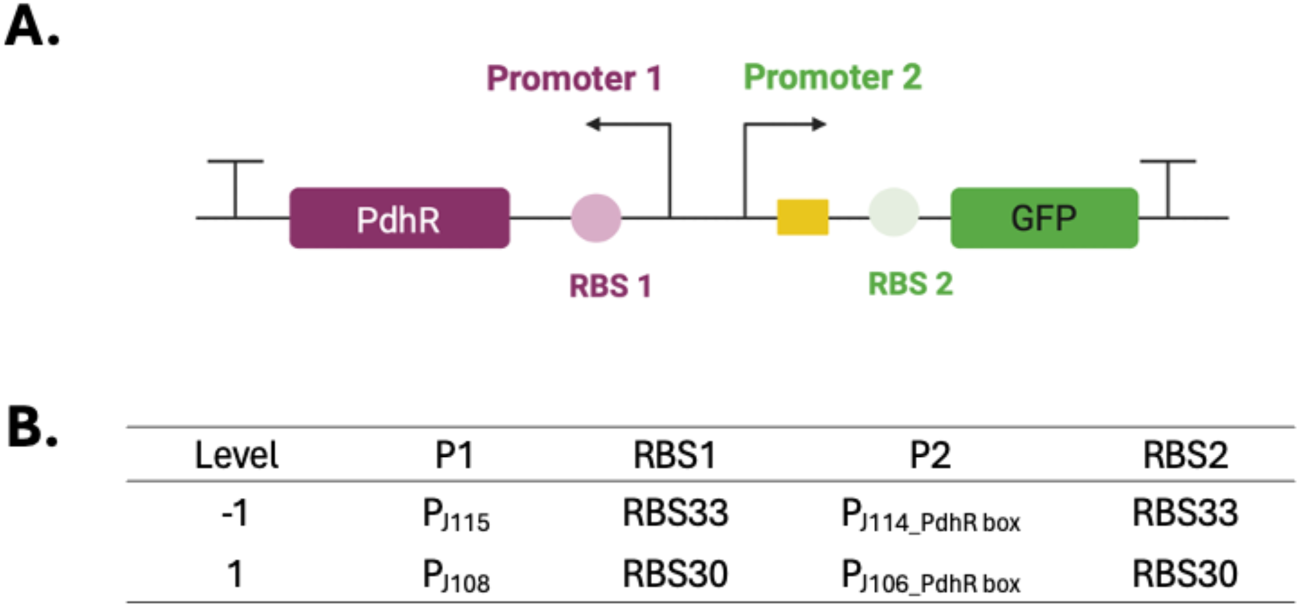
Genetic configuration of biosensor designs conforming to fractional factorial design. (A) The genetic circuits are shown, and the full table can be found in Table 1. (B) Levels of four independent factors were used in the fractional factorial design. P1 and P2 refer to Promoter 1 and Promoter 2, respectively. Each factor was tested at two levels: 1 was a high level, while -1 represented a low level.

### 4.5 Quantification of pyruvate

To evaluate the sensing performance and establish the correlation between biosensor output and actual pyruvate level, both extracellular and intracellular pyruvate concentration was quantified. For extracellular pyruvate detection, samples were centrifuged at 8000 rpm for 5mins, and supernatant was measured by ICS-5000 high-performance liquid chromatography (HPLC) with an Aminex HPX-87H column. A 2 mM H_2_SO_4_ solution was used as eluent at a flow rate of 0.6 ml/min and column oven temperature of 60 °C. Pyruvate was detected on an ultraviolet (UV) detector at 210nm and quantified based on an external standard of sodium pyruvate. Intracellular pyruvate was analyzed by first breaking cells using a methanol-water solution and three freeze-thaw cycles^48^. Briefly, 1mL cells were harvested by centrifugation at 8000 rpm for 5 minutes, washed twice with cold 0.9% (w/v) sodium chloride to remove extracellular pyruvate, and resuspended in 1 mL of pre-chilled methanol-water (50:50, v/v) solution at -20 °C. Cells were disrupted by three freeze–thaw cycles (freezing in -70 °C freezer for 30 minutes and thawing on ice for 4 mins with 1 minute of vortexing). The lysate was centrifuged at 14000 rpm for 10 minutes, and the supernatant containing intracellular pyruvate was collected. After this process, methanol was evaporated in a vacuum centrifuge (Concentrator Plus, Eppendorf) at 45°C under V-AQ model for 2 hours. After evaporation, 10µl MilliQ water was added to dissolve samples thoroughly and subsequently analyzed using the pyruvate assay kit (MAK332) following manufacturer’s protocols. To account for biomass variation, intracellular pyruvate concentration was normalized by dry cell weight (DCW). DCW was estimated based on a previously established calibration curve correlating OD₆₀₀ to DCW (Figure S4).

### 4.6 Data Processing and Modeling

All measurements were performed in triplicate. For pairwise comparisons between two independent groups, t-test was performed using the T.TEST function in Microsoft Excel 16.80, and statistical significance is indicated as \**P* < 0.05, ** *P* <0.01, *** *P* <0.001.

Responses of the biosensors were obtained by fitting the experimental data to the Hill equation (Figure 1A):

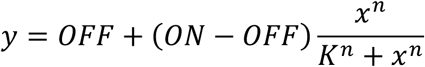

Where *y* is the relative fluorescence intensity (GFP/OD), *ON/OFF* is the maximum/ minimum relative fluorescence intensity, *x* is concentration of pyruvate, *K* is the threshold of the gene circuit and *n* is the Hill coefficient. Parameter estimation and Hill equation visualization were performed in GraphPad Prism 10.

The fractional factorial design form was designed by JMP Pro 17 software. Statistical data was performed by R version 4.3.2. The linear model was fit using the lm function, and the pareto plots were generated with the R paretoPlot functions^49^. The relationship between the responses and factors was expressed by the linear regression equation: 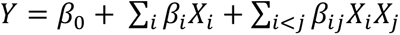.

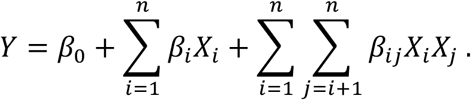

Where *Y* represents the predicted response, *X_i_* are the considered factors, *β_0_* is the interception coefficient, *β_i_* are the linear coefficient of the main effect (ME) and *β_ij_* are coefficients of the two-factor interactions (2FI). The quality of the regression equations was assessed according to the adjusted coefficient of determination (Adj R^2^). Statistical analysis of the model was performed using Analysis of Variance (ANOVA) and *P* < 0.05 was considered significant, and statistically significant is indicated as \**P* < 0.05, ** *P* <0.01, *** *P* <0.001.

The quality of the regression models was evaluated using the Akaike Information Criterion (AIC), which balances model fit and complexity. The Akaike equation (AIC formula) is as follows:

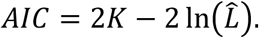

Where *K* represents the number of estimated parameters in the model and *L* is the maximized value of the likelihood function. AIC values calculated using the AIC function in R version 4.3.2.

For pairwise comparisons between two groups, an unpaired two-tailed Welch’s t-test was used to evaluate statistical significance, assuming unequal variances.

The schematic illustrations were created with BioRender (BioRender.com).

## 5. Author information

### Corresponding Authors

Pieter Candry – Laboratory of Systems and Synthetic Biology, Wageningen University & Research, Stippeneng 4, 6708 WE, Wageningen, The Netherlands

### Authors

Zihan Gao – Laboratory of Systems and Synthetic Biology, Wageningen University & Research, Stippeneng 4, 6708 WE, Wageningen, The Netherlands

Maria Suarez-Diez – Laboratory of Systems and Synthetic Biology, Wageningen University & Research, Stippeneng 4, 6708 WE, Wageningen, The Netherlands

## Author contributions

Z.G. performed the experiments, analyzed the results, and wrote the initial draft. P.C. and M.S.D. guided the experimental design and data analysis. All authors contributed to writing and reviewing the manuscript.

## Notes

The authors declare no competing financial interest.

## Acknowledgements

The authors acknowledge the assistance of Dr. Zeynep Efsun Duman-Özdamar for valuable guidance of data analysis, Dr. Enrique Asin-Garcia for strain selection and helpful discussions and Silvia Rodríguez Marcos for experiment material support. Zihan Gao was supported by the China Scholarship Council (CSC) under Grant No.202304910065. Pieter Candry was supported by the Dutch Research Council (NWO) Veni Talent Programme (File no. 21027) and the Dutch Sectorplan Bèta-II - Biology. Fig. 1, Fig.2 and Fig.3A were created with BioRender (BioRender.com). Fig. 1C fractional factorial design matrix adapted from Kevin Dunn’s DoE teaching materials (Dunn, 2025).

## 6. Data availability statement

Data will be made available upon reasonable request.

