## Supplemental Information for "Design-Build-Test-Learn guided engineering of a whole-cell pyruvate biosensor based on transcription factor"

MANUSCRIPT

**Supplementary material**

**Table S1 Strains used in this study**

| Strain | Description | Description |
|---|---|---|
| *E.coli* DH5α | Wild type | Lab stored |
| *E.coli* MG1655 | Wild type | Lab stored |
| *E.coli* K-12 BW25113△pdhR | pdhR knock out strain, kana resistance | KEIO collection |
| No. 0 | *E.coli* △pdhR containing 0_ pUC19_$P_{J115}$_RBS30_PdhR _$P_{pdh}$_ PdhR box_ RBS30_sfGFP | This study |
| △pdhR pUC19_sfGFP | *E.coli* △pdhR containing pUC19_ $P_{J106}$_PdhR box_RBS30_sfGFP | This study |
| △pdhR pUC19_ PdhR_sfGFP | *E.coli* △pdhR containing pUC19_ $P_{J115}$_RBS30_PdhR_$P_{J106}$_ PdhR box _RBS30_sfGFP | This study |
| No. 1 | *E.coli* △pdhR containing 1_pUC19_$P_{J115}$_RBS33_PdhR _$P_{J114}$_ PdhR box_RBS33_sfGFP | This study |
| No. 2 | *E.coli* △pdhR containing 2_pUC19_$P_{J115}$_RBS30_PdhR _$P_{J114}$_ PdhR box_RBS30_sfGFP | This study |
| No. 3 | *E.coli* △pdhR containing 3_pUC19_$P_{J115}$_RBS33_PdhR _$P_{J106}$_ PdhR box_RBS30_sfGFP | This study |
| No. 4 | *E.coli* △pdhR containing 4_pUC19_$P_{J115}$_RBS30_PdhR _$P_{J106}$_ PdhR box_RBS33_sfGFP | This study |
| No. 5 | *E.coli* △pdhR containing 5_pUC19_$P_{J108}$_RBS33_PdhR _$P_{J114}$_ PdhR box_RBS30_sfGFP | This study |
| No. 6 | *E.coli* △pdhR containing 6_pUC19_$P_{J108}$_RBS30_PdhR _$P_{J114}$_ PdhR box_RBS33_sfGFP | This study |
| No. 7 | *E.coli* △pdhR containing 7_pUC19_$P_{J108}$_RBS33_PdhR _$P_{J106}$_ PdhR box_RBS33_sfGFP | This study |
| No. 8 | *E.coli* △pdhR containing 8_pUC19_$P_{J108}$_RBS30_PdhR_$P_{J106}$_ PdhR box_RBS30_sfGFP | This study |

**Table S2 Plasmids used in this study**

| Plasmid | Description | Description |
|---|---|---|
| pUC19 | Amp, ColE1 oign, high copy number | Lab stored |
| 0_pUC19_PdhR_Ppdh_sfGFP | 0_ pUC19_P$_{J115}$_RBS30_PdhR _P$_{pdh}$_ PdhR box_ RBS30_sfGFP | This study |
| pUC19_sfGFP | pUC19_ P$_{J106}$_PdhR box_RBS30_sfGFP | This study |
| pUC19_ PdhR_sfGFP | pUC19_ P$_{J115}$_RBS30_PdhR_P$_{J106}$_ PdhR box _RBS30_sfGFP | This study |
| 1_pUC19_PdhR_sfGFP | 1_pUC19_P$_{J115}$_RBS33_PdhR _P$_{J114}$_ PdhR box_RBS33_sfGFP | This study |
| 2_pUC19_PdhR_sfGFP | 2_pUC19_P$_{J115}$_RBS30_PdhR _P$_{J114}$_ PdhR box_RBS30_sfGFP | This study |
| 3_pUC19_PdhR_sfGFP | 3_pUC19_P$_{J115}$_RBS33_PdhR _P$_{J106}$_ PdhR box_RBS30_sfGFP | This study |
| 4_pUC19_PdhR_sfGFP | 4_pUC19_P$_{J115}$_RBS30_PdhR _P$_{J106}$_ PdhR box_RBS33_sfGFP | This study |
| 5_pUC19_PdhR_sfGFP | 5_pUC19_P$_{J108}$_RBS33_PdhR _P$_{J114}$_ PdhR box_RBS30_sfGFP | This study |
| 6_pUC19_PdhR_sfGFP | 6_pUC19_P$_{J108}$_RBS30_PdhR _P$_{J114}$_ PdhR box_RBS33_sfGFP | This study |
| 7_pUC19_PdhR_sfGFP | 7_pUC19_P$_{J108}$_RBS33_PdhR _P$_{J106}$_ PdhR box_RBS33_sfGFP | This study |
| 8_pUC19_PdhR_sfGFP | 8_pUC19_P$_{J108}$_RBS30_PdhR_P$_{J106}$_ PdhR box_RBS30_sfGFP | This study |

**Table S3 List of genetic parts and sequences used in this study**

Underlined sequences indicate -35 and -10 prompter region. Sequences in red are PdhR binding sites.

| Part name | Type and source | DNA sequence (5'-3') |
|---|---|---|
| $P_{J115}$ (J23115) | Constitutive promoter[1] | TTTATAGCTAGCTCAGCCCTTGGTACAATGCTAGC |
| $P_{J114}$ (J23114) | Constitutive promoter | TTTATGGCTAGCTCAGTCCTAGGTACAATGCTAGC |
| $P_{J108}$ (J23108) | Constitutive promoter | CTGACAGCTAGCTCAGTCCTAGGTATAATGCTAGC |
| $P_{J106}$ (J23106) | Constitutive promoter | TTTACGGCTAGCTCAGTCCTAGGTATAGTGCTAGC |
| RBS30 (B0030) | Ribosome binding site[2] | ATTAAAGAGGAGAAA |
| RBS33 (B0033) | Ribosome binding site | TCACACAGGAC |
| Ppdh | Inducible promoter | TGGACATAAGGTGAATACTTTGTTACTTTAGCGTCACAGACATGAAATTGGTAAGACCAATTGACTTCGGCAAGTGGCTTAAGACAGGAACTC |
| PdhR | Transcription factor (Amplified from *E.coli* MG1655 by PCR) | ATGGCCTACAGCAAAATCCGCCAACCAAAACTCTCCGATGTGATTGAGCAGCAACTGGAGTTTTTGATCCTCGAAGGCACTCTCCGCCCGGGCGAAAAACTCCCACCGGAACGCGAACTGGCAAAACAGTTTGACGTCTCCCGTCCCTCCTTGCGTGAGGCGATTCAACGTCTCGAAGCGAAGGGCTTGTTGCTTCGTCGCCAGGGTGGCGGCACTTTTGTCCAGAGCAGCCTATGGCAAAGCTTCAGCGATCCGCTGGTGGAGCTGCTCTCCGACCATCCTGAGTCACAGTATGACTTGCTCGAAACACGACACGCCCTGGAAGGTATCGCCGCTTATTACGCCGCGCTGCGTAGTACCGATGAAGACAAGGAACGCATCCGTGAACTCCACCACGCCATAGAGCTGGCGCAGCAGTCTGGCGATCTGGACGCGGAATCAAACGCCGTACTCCAGTATCAGATTGCCGTCACCGAAGCGGCCCACAATGTGGTTCTGCTTCATCTGCTAAGGTGTATGGAGCCGATGTTGGCCCAGAATGTCCGCCAGAACTTCGAATTGCTCTATTCGCGTCGCGAGATGCTGCCGCTGGTGAGTAGTCACCGCACCCGCATATTTGAAGCGATTATGGCCGGTAAGCCGGAAGAAGCGCGCGAAGCATCGCATCGCCATCTGGCCTTTATCGAAGAAATTTTGCTCGACAGAAGTCGTGAAGAGAGCCGCCGTGAGCGTTCTCTGCGTCGTCTGGAGCAACGAAAGAATTAG |

| | | |
|---|---|---|
| sfGFP | Report gene | ATGCGTAAAGGCGAAGAGCTGTTCACGGGCGTAGTTCCGATTCTGGTCGAGCTGGACGGCGATGTGAACGGTCATAAGTTTAGCGTTCGCGGTGAAGGTGAGGGCGACGCGACCAACGGCAAACTGACCCTGAAGTTCATCTGCACCACCGGTAAACTGCCGGTGCCTTGGCCGACCTTGGTGACGACGTTGACGTATGGCGTGCAGTGTTTTGCGCGTTATCCGGACCACATGAAACAACACGATTTCTTCAAATCTGCGATGCCGGAGGGTTACGTCCAGGAGCGTACCATTTCCTTCAAGGATGATGGCTACTACAAAACTCGCGCAGAGGTTAAGTTTGAAGGTGACACGCTGGTCAATCGTATCGAATTGAAGGGTATCGACTTTAAAGAGGATGGTAACATTCTGGGCCATAAACTGGAGTATAACTTCAACAGCCATAATGTTTACATTACGGCAGACAAGCAAAAGAACGGCATCAAGGCCAATTTCAAGATTCGCCACAATGTTGAGGACGGTAGCGTCCAACTGGCCGACCATTACCAGCAGAACACCCCAATTGGTGACGGTCCGGTTTTGCTGCCGGATAATCACTATCTGAGCACCCAAAGCGTGCTGAGCAAAGATCCGAACGAAAAACGTGATCACATGGTCCTGCTGGAATTTGTGACCGCTGCGGGCATCACCCACGGTATGGATGAACTGTACAAA |

1: https://parts.igem.org/Part:BBa_J23115
2: https://parts.igem.org/Part:BBa_B0030

**Table S4 Primer used in this study**

| Primer name | DNA sequence (5'-3') | Usage |
|---|---|---|
| pUC_rev | GAACGCTCTCATCCCCGGGTACCGAGCTC | For ampliying backbone of plasmid for assembling gene into plasmid |
| pUC_fwd | TAGCTATAAACCTCTAGAGTCGACCTGCAGGCATGCAAGCT | |
| PdhR_T_fwd | ACTCTAGAGGTTTATAGCTAGCTCAGCCCTTGGTACAATGCTAGCAGGTCGACATTAAAGAGGAGAAATCTAGAGGATGGCCTACAGCAAAATCCG | For amplify *pdhR* gene and adding promoter and RBS |
| PdhR_T_rev | ACCCGGGGATGAGAGCGTTCACCGACAAACAACAGATAAAACGAAAGGCCCAGTCTTTCGACTGAGCCTTTCGTTTTATTTGATGCCTGGCTAATTCTTTCGTTGCTCCAGACGAC | |
| pUC2_fwd | GAACGCTCTCAGAGTCGACCTGCAGGCATG | For ampliying backbone of plasmid |
| pUC2_rev | TAGCCGTAAAAGAGGTTTATAGCTAGCTCAGCCCTTG | |
| sfGFP_T_fwd | ATAAACCTCTTTTACGGCTAGCTCAGTCCTAGGTATAGTGCTAGCGTCATGCAAGCATTGGTAAGACCAATACGCCTATTAAAGAGGAGAAAGCCAACGTATGCGTAAAGGCGAAGAGCTG | For amplify *sfGFP* gene and adding promoter, RBS and terminator |
| sfGFP_T_rev | GGTCGACTCTGAGAGCGTTCACCGACAAACAACAGATAAAACGAAAGGCCCAGTCTTTCGACTGAGCCTTTCGTTTTATTTGATGCCTGGTCATTTGTACAGTTCATCCATACCATGC | |
| verify_PG _fwd | GTTGTGTGGAATTGTGAGCGG | For verifying *PdhR*, s*fGFP* and various promoters and RBS |
| verify_PG_rev | TCCCAGTCACGACGTTGTAAAAC | |
| verify_pUC19_1 | TACATCGAACTGGATCTCAACAG | For verifying plasmid |
| verify_pUC19_2 | GATCCTTTTTGATAATCTCATGACC | |
| Ppdh_fwd | TAATAGGCGTGAGTTCCTGTCTTAAGCC | For amplifying $P_{pdh}$ from *E.coli* genome |
| Ppdh_rev | ATAAACCTCTGTCTACATTTGTGCATAGTTAC | |
| backbone_fwd | AAATGTAGACAGAGGTTTATAGCTAGCTCAG | For amplying backbone and assembling $P_{pdh}$ into plasmid |
| backbone_rev | ACAGGAACTCACGCCTATTAAAGAGGAG | |
| pUC19_Ppdh_fwd | TCTAGAGGATGGCCTACAGC | For amplifying backbone of plasmid containg *pdhR* and |
| pUC19_Ppdh_rev | GCCAACGTATGCGTAAAGGC | |

| | | |
|---|---|---|
| | | *sfGFP* gene for assembing new promoters and RBS into plasmid |
| verify_PdhR_fwd | GTTAAAATGTGCACAGTTTCATG | For verifying the absence of *pdhR* gene in *E.coli* ΔpdhR genome |
| verify_PdhR_rev | ATGCGCTTGATTTACAACATC | |

**Table S5 Best fits for the responses of various biosensors in this study**

| No. | P1 | RBS1 | P2 | RBS2 | n (Hillslope) | K ($EC_{50}$) | $R^2$ |
|---|---|---|---|---|---|---|---|
| 0 | -1 | 1 | n.a | 1 | 1.57 ± 0.10 | 1.25 ± 0.01 | 0.9930 ± 0.0024 |
| 1 | -1 | -1 | -1 | -1 | 1.79 ± 0.11 | 1.59 ± 0.08 | 0.9687 ± 0.0056 |
| 2 | -1 | 1 | -1 | 1 | 1.64 ± 0.14 | 0.98 ± 0.05 | 0.9784 ± 0.0059 |
| 3 | -1 | -1 | 1 | 1 | 1.98 ± 0.15 | 1.66 ± 0.07 | 0.9981 ± 0.0005 |
| 4 | -1 | 1 | 1 | -1 | 1.56 ± 0.18 | 1.27 ± 0.09 | 0.9778 ± 0.0011 |
| 5 | 1 | -1 | -1 | 1 | 1.56 ± 0.17 | 1.65 ± 0.09 | 0.9794 ± 0.0105 |
| 6 | 1 | 1 | -1 | -1 | 1.64 ± 0.06 | 2.29 ± 0.07 | 0.9873 ± 0.0026 |
| 7 | 1 | -1 | 1 | -1 | 1.88 ± 0.14 | 1.52 ± 0.05 | 0.9652 ± 0.0057 |
| 8 | 1 | 1 | 1 | 1 | 2.06 ± 0.08 | 1.4 ±0.05 | 0.9714 ± 0.0025 |

**Table S6 Statistics of regression equations**

Dependent variable: Span (ON-OFF)

| Term | Estimate | *P* value | Significance |
|---|---|---|---|
| Intercept | 8631.3 | 0.0000264 | *** |
| P1 | -345.6 | 0.1788 | |
| RBS1 | -1100.7 | 0.0114 | * |
| P2 | 321.8 | 0.2021 | |
| RBS2 | -554.0 | 0.0677 | |

*Y(Span)* = 8631.3 -345.6*P1*-1100.7*RBS1*+321.8*P2*-554.0*RBS2*

$R^2$ = 0.9369; Adjusted $R^2$ = 0.8528; p-value = 0.03811

Dependent variable: Dynamic range ($\log_{10}$)

| Term | Estimate | *P* value | Significance |
|---|---|---|---|
| Intercept | 0.862178 | 0.000858 | *** |
| P1 | 0.081294 | 0.289472 | |
| RBS1 | 0.007065 | 0.918226 | |
| P2 | 0.005274 | 0.938878 | |
| RBS2 | -0.428494 | 0.006599 | ** |

*Y(Dynamic range)* = 0.862178 +0.081294*P1*+0.007065*RBS1*+0.005274*P2*-0.428494*RBS2*

$R^2$ = 0.9405; Adjusted $R^2$ = 0.8612; p-value = 0.03497

Dependent variable: OFF ($\log_{10}$)

| Term | Estimate | *P* value | Significance |
|---|---|---|---|
| Intercept | 3.18898 | 0.0000282 | *** |
| P1 | -0.1191 | 0.20863 | |
| RBS1 | -0.04822 | 0.56403 | |
| P2 | -0.02036 | 0.8026 | |
| RBS2 | 0.49452 | 0.00699 | ** |

*Y(OFF)* = 3.18898-0.1191*P1*-0.04822*RBS1-0,02036P2+0.49452*

$R^2$ = 0.94; Adjusted $R^2$ = 0.86; p-value = 0.03543

Dependent variable: ON ($\log_{10}$)

| Term | Estimate | *P* value | Significance |
|---|---|---|---|
| Intercept | 4.05115 | 0.00000017 | *** |
| P1 | -0.03781 | 0.1158 | |
| RBS1 | -0.04116 | 0.0969 | |
| P2 | -0.01509 | 0.4457 | |
| RBS2 | 0.06602 | 0.0313 | * |

*Y(ON)* = 4.05115 -0.03781*P1*-0.04116*RBS1*-0.01509*P2*+0.06602*RBS2*

$R^2$ = 0.8964; Adjusted $R^2$ = 0.7583; p-value = 0.07814

Dependent variable: Dynamic range ($\log_{10}$)

| Term | Estimate | *P* value | Significance |
|---|---|---|---|
| Intercept | 0.862178 | 0.0017 | ** |
| P1 | 0.081294 | 0.1058 | |
| RBS1 | 0.007065 | 0.86117 | |
| P2 | 0.005274 | 0.89591 | |
| RBS2 | -0.428494 | 0.00635 | ** |
| P1:RBS1 | 0.097441 | 0.11178 | |

*Y(Dynamic range)* = 0.862178+0.081294*P1+0.007065RBS1+0.005274P2-*
0.428494*RBS2*+0.09744P1**RBS1*

$R^2$ = 0.9874; Adjusted $R^2$ = 0.9561; p-value = 0.03109

| Dependent variable: Dynamic range ($log_{10}$) | | | |
|---|---|---|---|
| Term | Estimate | *P* value | Significance |
| Intercept | 0.86218 | 0.00000462 | *** |
| P1 | 0.08129 | 0.0336 | * |
| RBS2 | -0.42849 | 0.0000745 | *** |
| P1:RBS1 | 0.09744 | 0.019 | * |
| *Y(Dynamic range)* = 0.86218+0.08129*P1*-0.42849*RBS2*+0.09744*P1*RBS1* | | | |
| $R^2$ = 0.9871; Adjusted $R^2$ = 0.9774; p-value = 0.0003125 | | | |

**Table S7 Model comparison based on Akaike Information Criterion (AIC)**

| Model | Formula | AIC |
|---|---|---|
| 1 | $Y = \beta_0 + \beta_1 \cdot P1 + \beta_2 \cdot RBS1 + \beta_3 \cdot P2 + \beta_4 \cdot RBS2$ | -0,6566 |
| 2 | $Y = \beta_0 + \beta_1 \cdot P1 + \beta_2 \cdot RBS1 + \beta_3 \cdot P2 + \beta_4 \cdot RBS2 + \beta_5 \cdot P1 \cdot RBS1$ | -11,1058 |
| 3 | $Y = \beta_0 + \beta_1 \cdot P1 + \beta_2 \cdot RBS1 + \beta_3 \cdot RBS2 + \beta_4 \cdot P1 \cdot RBS1$ | -13,0143 |
| 4 | $Y = \beta_0 + \beta_1 \cdot P1 + \beta_2 \cdot RBS2 + \beta_3 \cdot P1 \cdot RBS1$ | -14,8603 |
| 5 | $Y = \beta_0 + \beta_1 \cdot P1 + \beta_2 \cdot RBS2$ | -4,6051 |
| 6 | $Y = \beta_0 + \beta_1 \cdot RBS2$ | -3,1218 |

**Table S8 Best fits for the dose–response curves of biosensor response to intracellular pyruvate**

| OFF (a.u) | ON (a.u) | Span (ON-OFF) | n (Hillslope) | K ($EC_{50}$) | $R^2$ |
|---|---|---|---|---|---|
| 160.97 ± 29.06 | 9854.33 ± 181.49 | 9292.67 ± 203.08 | 2.62 ± 0.05 | 3.41 ±0.06 | 0.9892 ± 0.0019 |

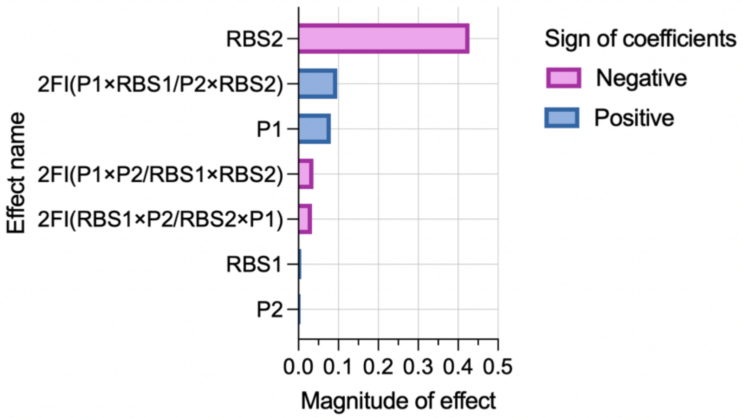


**Figure S1 Linear regression analysis of all interactions between factors**

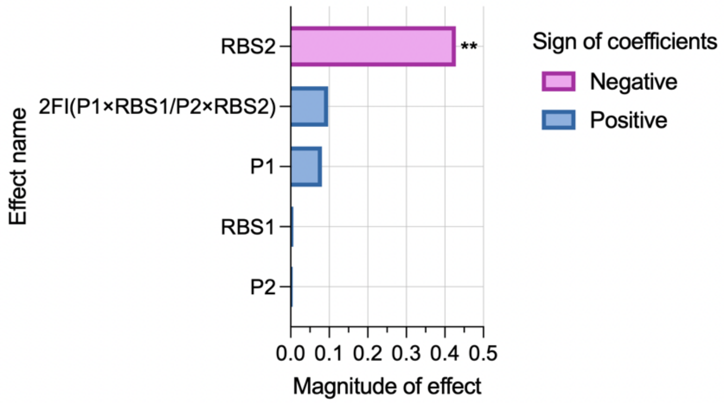


**Figure S2 Linear regression analysis of four main factors and two-factor interaction between P1×RBS1/P2×RBS2**

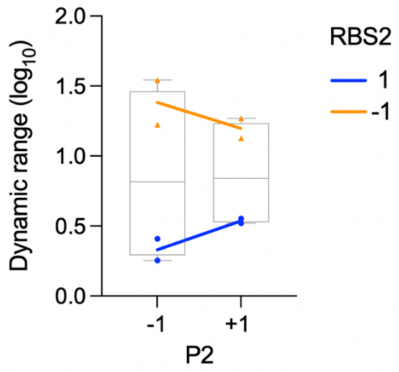


**Figure S3 Interaction effects between P2 and RBS2.** Blue circles and orange triangles indicate data points under high (1) and low (-1) levels of RBS2, respectively. The lines connect group means and illustrate the interaction effect: under high RBS2 (blue line), whereas under low RBS2 (orange line).

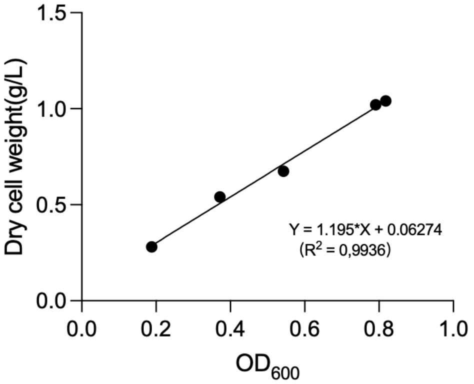


**Figure S4 Calibration curve correlating $OD_{600}$ to DCW**
